# Emergence of an invariant representation of texture in primate somatosensory cortex

**DOI:** 10.1101/646042

**Authors:** Justin D. Lieber, Sliman J. Bensmaia

## Abstract

A major function of sensory processing is to achieve neural representations of objects that are stable across changes in context and perspective. Small changes in exploratory behavior can lead to large changes in signals at the sensory periphery, thus resulting in ambiguous neural representations of objects. Overcoming this ambiguity is a hallmark of human object recognition across sensory modalities. Here, we investigate how the perception of tactile texture remains stable across exploratory movements of the hand, including changes in scanning speed, despite the concomitant changes in afferent responses. To this end, we scanned a wide range of everyday textures across the fingertips of Rhesus macaques at multiple speeds and recorded the responses evoked in tactile nerve fibers and somatosensory cortical neurons. We found that individual cortical neurons exhibit a wider range of speed-sensitivities than do nerve fibers. The resulting representations of speed and texture in cortex are more independent than are their counterparts in the nerve and account for speed-invariant perception of texture. We demonstrate that this separation of speed and texture information is a natural consequence of previously described cortical computations.

## INTRODUCTION

We are endowed with a remarkable ability to identify objects across a wide range of contexts and perspectives. For example, we can visually identify objects in a fraction of a second, even over broad changes in lighting, distance, or viewing angle. Similarly, we can auditorily identify the timbre of voices and musical instruments across a wide range of loudness and pitches (Handel and Erickson 2001; Marozeau et al. 2003). In both vision and audition, these perceptual invariances are achieved despite sensory representations at the periphery (the retina, the cochlea) that are highly dependent on perspective and context (Enroth-Cugell and Robson 1966; Sachs and Young 1979; Croner and Kaplan 1995; Joris et al. 2011). Indeed, a signature of sensory processing is a progression of object representations that become increasingly robust to changes in context (Avidan et al. 2002; Finn et al. 2007; Sadagopan and Wang 2008; Walker et al. 2011; Cadieu et al. 2014; Metzen et al. 2016).

In touch, the best known instance of perceptual invariance is for texture: tactile texture perception has been shown to be nearly independent of the force exerted on the surface (Lederman and Taylor 1972; Lederman 1981) or the speed at which it is scanned across the skin (Lederman 1974; Meftah et al. 2000; Boundy-Singer et al. 2017). Remarkably, this perceptual invariance is achieved despite responses in the somatosensory nerves that are strongly modulated by exploratory parameters such as scanning speed (Goodwin and Morley 1987a; Phillips et al. 1992; DiCarlo and Johnson 1999) and, to a lesser degree, force (Goodwin and Morley 1987b; Phillips et al. 1992; Saal et al. 2018). The effect of scanning speed on texture coding in the nerve is particularly pronounced for fine textures, which are encoded in precisely timed, texture-specific spiking sequences that contract or dilate multiplicatively with increases and decreases in speed, respectively (Weber et al. 2013). Thus, to achieve an invariant percept of texture, texture-specific information must be extracted from peripheral signals that are highly dependent on exploratory parameters.

As texture representations ascend the somatosensory neuraxis towards somatosensory cortex, precisely timed patterns of spatio-temporal activity are processed by canonical sensory transformations, such as differentiating filters that calculate spatial (DiCarlo and Johnson 2000; Sripati et al. 2006; Bensmaia et al. 2008) and temporal (DiCarlo and Johnson 2000; Sripati et al. 2006; Saal et al. 2015) variation across the peripheral signal. These filters extract perceptually-relevant stimulus information that may not be present in the firing rates of peripheral afferents (Connor 1990). It thus stands to reason that these same mechanisms could extract a speed-invariant representation of texture that was not present in the peripheral response. Indeed, previous work suggests that a subpopulation of neurons in somatosensory cortex may exhibit speed-invariant responses to texture (Sinclair and Burton 1991; DiCarlo and Johnson 1999; Dépeault et al. 2013; Bourgeon et al. 2016). However, these studies only characterized cortical responses to parametrically defined dot patterns and gratings that span a narrow range of tangible textures (Weber et al. 2013). Furthermore, these studies focused primarily on the speed-sensitivity of cortical neurons without comparing these effects to those seen in peripheral afferents.

In the present study, we seek to fill this gap by recording the responses of neurons in somatosensory cortex – including Brodmann’s areas 3b, 1, and 2 – to natural textures scanned over the skin at various speeds, spanning the range used in natural texture exploration (Morley et al. 1983; Gamzu and Ahissar 2001; Libouton et al. 2010; Tanaka et al. 2014; Callier et al. 2015). Using these data, we then directly compare how neuronal firing rate responses are modulated by speed in the nerve and in cortex. We find that while speed modulation is generally weaker in cortical firing rates than in afferent firing rates, this effect does not typically confer a speed-invariant texture code to individual cortical neurons. Rather, we find that speed-invariant texture perception is best explained by an untangling of information about speed and texture across the responses of neuronal populations. The resulting cortical population response better accounts for speed-invariant texture perception than does its peripheral counterpart.

## METHODS

### Experimental Methods

#### Peripheral texture responses

We recorded the responses of 39 tactile fibers in six anesthetized macaque monkeys to 55 textured surfaces scanned over the fingertip, including everyday textures such as fabrics, sandpapers, as well as plastic gratings and embossed dots. Recordings were collected from afferents innervating the distal fingertip, using standard methods (Talbot et al. 1968). For 21 of these 39 afferents (9 slowly adapting type 1 – SA1 – fibers, 9 rapidly-adapting – RA – fibers, and 3 Pacinian – PC – fibers), we were able to maintain isolation long enough to record responses at 3 different scanning speeds, namely 40, 80, and 120 mm/s (all ± 0.1 mm/s, typically for 1, 2, and 4 repetitions at each speed, respectively). In these experiments, texture presentation was blocked by speed rather than by texture. That is, we first recorded the response of afferents to all 55 textures at 80 mm/s, then at 40 mm/s or 120 mm/s, and in the third block at the remaining speed. Our analyses only consider responses during the steady-state contact period for force and speed, which lasted for at 2, 1, and 0.5 seconds, at 40, 80, and 120 mm/s, respectively. For 14/21 afferents, we had only 1 repetition at 40 mm/s. We confirmed with our remaining 7 afferents (2 SA1s, 2 PCs, 3 RAs) that using single trials to calculate the firing rate at 40 mm/s introduced minimal error to the estimation of speed-sensitivity (median absolute error between subsamples < 4% / doubling). Anesthesia was maintained using isoflurane. All experimental procedures complied with the National Institutes of Health Guide for the Care and Use of Laboratory Animals and were approved by the Animal Care and Use Committee of the University of Chicago. The neurophysiological approach is described in more detail in previously published articles describing studies that used a subset of these data (Weber et al. 2013; Lieber et al. 2017).

#### Cortical texture responses

Extracellular recordings were made in the postcentral gyri of three hemispheres in each of three awake macaque monkeys (all male, 6-8 yrs old, 8-11 kg). On each recording day, a multielectrode microdrive (NAN Instruments, Nazaret Illit, Israel) was loaded with three electrodes (tungsten, Epoxylite insulated, FHC Inc.), spaced 650-μm apart, which were driven into cortex until they encountered neurons from Brodmann’s areas 3b, 1, and 2 of with RFs on the distal fingerpad of digits 2-5. The respective cortical fields were identified based on anatomical landmarks, RF location, and response properties. Recordings were obtained from neurons that met the following criteria: (1) action potentials were well isolated from the background noise, (2) the finger could be positioned such that the textured surface impinged on the center of the RF, and (3) the neuron was clearly driven by light cutaneous touch. Isolations had to be maintained for at least 30 minutes to complete 5 repetitions of the basic texture protocol (59 different textures presented at 80 mm/s) and an additional 25 minutes to complete 5 repetitions of the speed protocol (10 different textures presented at 4 different speeds).

Responses from 141 single units were obtained for the basic texture protocol: 35 units from area 3b, 81 units from area 1, and 25 units from area 2. For 49 of these single units, we were able to maintain isolation long enough to obtain responses for the speed protocol: 14 units from area 3b, 26 units from area 1, and 9 units from area 2. In this protocol, ten textures were presented at 4 different speeds: 60, 80, 100, and 120 mm/s (all ± 1.1 mm/s). We opted not to test textures at 40 mm/s as we had in the peripheral nerve experiments because we subsequently discovered that this is well below the typical range of speeds used to explore textures (Callier et al. 2015). Four of the textures (satin, chiffon, nylon, and hucktowel) were also used in our recordings of afferent responses. The other six textures were chosen to contain an overlay of both fine (< 1 mm) and coarse (> 1 mm) spatial features (fabric grating [wide spacing], sunbrella upholstery, fuzzy upholstery, faux croc skin, 7.7 mm dots, 7.7 mm dots / 1 mm grating overlay). Each texture was presented 5 times at each speed, for 2.3, 1.7, 1.4, and 1.2 seconds at 60, 80, 100, and 120 mm/s, respectively. Textures and speeds were presented in pseudo-random order. All experimental procedures complied with the National Institutes of Health Guide for the Care and Use of Laboratory Animals and were approved by the Animal Care and Use Committee of the University of Chicago. The neurophysiological approach is described in more detail in a previously published article describing a study that used a subset of these data (Lieber and Bensmaia 2019).

#### Roughness magnitude estimation

Six subjects (5m, 1f, ages 18-24) were passively presented with each of 59 textures presented at 80 mm/s and produced a rating proportional to its perceived roughness. This procedure was repeated 6 times over 6 blocks. Ratings were normalized within block and then averaged within subject. Because ratings were consistent across subjects (correlation: 0.87 ± 0.079), ratings were then averaged across subjects. All procedures were approved by the Institutional Review Board of the University of Chicago and all subjects provided informed consent. The psychophysical procedure has been previously described in detail in a previously published article describing a study using these data (Lieber and Bensmaia 2019).

### Analysis

#### Firing rates and speed effects

Peripheral firing rates were calculated over windows of 2, 1, and 0.5 seconds for textures presented at 40, 80, and 120 mm/s, respectively. Cortical firing rates were calculated over windows of 2.3, 1.7, 1.4, and 1.2 seconds for textures presented at 60, 80, 100, and 120 mm/s, respectively. All cortical firing rates were corrected for baseline firing. For each neuron, the firing rate over a 500-ms period before each trial was averaged over all trials to obtain the baseline firing rate, which was then subtracted from the neuron’s texture-evoked firing rates.

To minimize any systematic biases due to differences in texture sets across the peripheral and cortical experiments, we selected textures in the peripheral experiment to match the 10 textures used in the cortical experiment. As four textures were used in both experiments, we sought appropriate matches for the remaining six. To this end, we examined the subset of 20 textures for which we had cortical responses at 80 mm/s and peripheral responses at all three speeds. To assess similarity between the speed-set of 6 textures and the shared set of 20 textures, we calculated the Euclidian distance between texture pairs based on trial-averaged firing rates across the cortical population (6*20=120 pairs) and then selected the 6 pairs with the shortest distance. That is, we chose the 6 textures in the shared set of 20 textures that evoked the most similar pattern of responses across the cortical population to the 6 textures in the speed set. We then used these 6 textures to round out the peripheral set. The underlying assumption is that if two textures evoke similar patterns of responses across cortical neurons, they would also evoke similar responses across peripheral afferents. For all analyses in the manuscript, we achieved qualitatively similar results when we used peripheral responses to the full set of 55 textures or to the 4 shared textures.

To quantify the effect of speed on texture-driven firing rates, we regressed firing rate on the log (base 2) of speed, having established that a logarithmic function better captures the effect of speed on firing rate than does a linear one (cf. (Essick and Edin 1995; DiCarlo and Johnson 1999)).

#### Calculating speed-sensitivity, speed/texture ratio, and response heterogeneity

We wished to quantify the relative effect of speed and texture on each afferent and neuron. To this end, we calculated three quantities for each cell. First, we defined speed-sensitivity as the slope of the linear regression between log speed and firing rate, normalized by the mean firing rate at 80 mm/s evoked by all 10 textures (to correct for overall firing rate differences across neurons). Second, we defined texture-sensitivity as the across-texture coefficient of variation: that is, the standard deviation across the firing rate responses to a set of 24 textures used in both experiments, (presented at 80 mm/s, see (Lieber and Bensmaia 2019)), normalized by the average firing rate of those same 24 textures. Finally, we defined the speed/texture ratio as the ratio of speed-sensitivity to texture-sensitivity.

Because this ratio metric becomes unstable if texture-sensitivity is near zero, we verified that all neurons exhibited sufficiently large values of texture-sensitivity (all peripheral afferents > 0.37, all cortical neurons > 0.28). Note further that, for all neurons, texture-sensitivity was significantly greater than that expected by chance, based on a permutation test that compares the measured texture-sensitivity to that obtained when single-trial responses are shuffled across texture labels (*p* < 0.05). We report that the cortical population contains a significant proportion of neurons with a smaller speed/texture ratio than any peripheral afferent. To test the reliability of this effect, we randomly shuffled the cortex/periphery labels on the combined population of afferents and cortical neurons, and recomputed the number of “neurons” with smaller speed/texture ratios than “afferents.” From this simulation, we could characterize the relative distribution of speed/texture ratios expected by chance.

#### Neural population analyses

We sought to evaluate the extent to which representations of texture and speed in the neural populations were either overlapping or well-separated. This required a three step process: 1) identify the texture representation in each population, 2) identify the speed-representation in each population, and 3) determine the amount of overlap between the two representations.

To identify the major axes of each population’s texture response, we applied a principal components analysis (PCA) to peripheral and cortical population responses, using only the responses to the set of 24 textures (presented at 80 mm/s) shared between the two data sets. This allowed us to identify the first D dimensions in each population that captured the majority of the texture response variance, which we refer to as *P*_*t*_. To identify the main speed-related axis in each population, we applied demixed principal components analysis (dPCA)(Kobak et al. 2016) to the full set of trial-averaged responses to textures presented at multiple speeds (periphery: 10 textures at 3 speeds, cortex: 10 textures at 4 speeds). Specifically, we first created a speed-marginalization of the population response by subtracting out each texture’s average firing rate (across all speeds) from the full response matrix. Thus, the full response matrix can be expressed as:

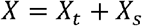

where *X* is the full texture response, *X*_*t*_ is the texture marginalization, and *X*_*s*_ is the speed marginalization (which, in the terminology of (Kobak et al. 2016), contains both the pure speed marginalization and speed-texture interaction marginalization). Next, we found the best linear mapping from the full texture response to the speed marginalization using least squares regression:

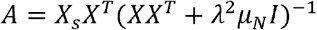

where *λ* is a regularization parameter (set to 10^−6^, as in (Kobak et al. 2016)) and *μ*_*N*_ is each neuron’s variance across all speed and texture conditions. Finally, we used PCA on the best linear approximation of the speed marginalization, *AX*, to find its primary axis of variation, which we refer to as *P*_*s*_. In the terminology of (Kobak et al. 2016), this corresponds to the primary encoder axis for the speed marginalization.

We validated each population’s primary speed-axis using two metrics. First, we confirmed that the projection of any given population response onto the speed axis covaried with the actual speed (in log units) by measuring the coefficient of determination (R^2^) between the two variables (periphery: 10×3=30 conditions, cortex: 10×4=40 conditions). Second, we quantified the extent to which the primary speed-axis accounted for speed-driven neural responses by computing the ratio of the variance captured by the single speed axis to the total variance in the speed marginalization (summed across neurons).

To determine the amount of overlap between the texture space and the speed dimension, we calculated an alignment index (Elsayed et al. 2016; Gallego et al. 2018), defined as the amount of speed-driven variance captured by the texture space, normalized by the total amount of speed-related variance in the population response. Specifically, we define the alignment index as:

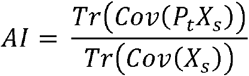

where *P*_*t*_ is defined as the principal axes of the texture space. We emphasize that, by construction, this metric is insensitive to the raw magnitude of speed-related fluctuations in the population response. We verified with simulations that when the speed/texture ratio of the population response is doubled or halved the alignment index stays constant. In this sense, the alignment index is more similar to a relative angle between to representations than to a measurement of raw speed-sensitivity. For Figure *2*, we recomputed the texture-space, speed-axis, and alignment index for different subsamples of 21 neurons within the cortical space (to control for systematic biases due to sample size), for either individual dimensions of *P*_*t*_ (Figure *2*A) and for a range of the first D dimensions of *P*_*t*_ (Figure *2*B).

#### Texture discriminability across changes in speed

We sought to quantify how well populations of neurons could discriminate pairs of textures, and to what extent discrimination was impaired by changes in speed. For an individual neuron responding to any pair of textures and pair of speeds, we define a signal-to-noise ratio (SNR):

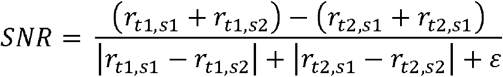

where *r*_*t*1,*s*1_ is the response (firing rate) of the neuron to texture *t*_1_ at speed *s*_1_. We added *ε* (= 1) to the denominator to avoid the instability in SNR caused by neurons that do not respond under some stimulus conditions. To compute the SNR for populations of neurons, we first collapsed the multidimensional, population-wide firing rates to a single discriminant value. To this end, we defined a population vector of firing rates *R*_*t*1,*s*1_ for a given texture *t*_1_ scanned at speed *s*_1_. We then found the line connecting the two textures’ average population response: (*R*_*t*1,*s*1_ + *R*_*t*1,*s*2_)−(*R*_*t*2,*s*1_ + *R*_*t*2,*s*2_). This line defines the texture-relevant axis in neural space. Next, we found the projections of the four population vectors onto this line, resulting in four scalar values, each corresponding to a texture, speed pair. To the extent that speed has no impact along the texture axis, the projections onto this line of the response vectors to each texture at different speeds will be identical. Finally, we computed the SNR on these projections in place of the full population vectors of rates.

#### Predicting roughness from neural responses

We sought to evaluate how well neural population responses could predict human judgments of surface roughness (55/59 textures for periphery and cortex, respectively, all presented at 80 mm/s). To this end, we implemented three distinct models: a mean firing rate regression model, a multiple regression model, and a second multiple regression model, constrained to be speed-invariant. The mean firing rate model comprised a single regressor, the mean firing rate across the full population of neurons (21 afferents, 49 cortical neurons). The multiple regression model included the first N principal components of the texture representation. For Figure 5, we chose a number of principal components where regression performance began to saturate (N=5), though the results were stable over different numbers of components (*Supplementary Figure 5*). For the constrained multiple regression model, we first found the primary speed axis in each population using dPCA (described above). Next, we removed the speed-axis projection from the full set of population firing rates. Finally, we performed the multiple regression as described before. This methodology ensured that the final regression weights were orthogonal to the primary speed axis and that the roughness predictions of the model were almost entirely speed-invariant.

For all three models, we used leave-one-out cross-validation to compute an equivalent of the coefficient of determination (*R*^2^). Specifically, for each texture in the set, we first fit the model using the other 23 texture responses as training data. We then applied that model to produce a prediction of roughness magnitude 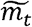 for the final, left-out texture. Across all textures, we compute the coefficient of determination as:

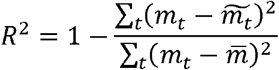

where *m*_*t*_ is the reported roughness.

To test the speed-invariance of these models, we applied each model (now fit on the full set of 55/59 textures) to texture responses at multiple speeds (10 textures in each set). We computed speed-sensitivity of the roughness predictions in a manner similar to that used for the neural data. First, we compute the slope of roughness (averaged across textures) vs. log_2_(speed). Then, we normalized the slopes by the average predicted roughness magnitude at 80 mm/s. As with the neuronal data, then, we report the speed-sensitivity of the roughness predictions as a percentage increase per doubling of speed.

#### Modeling neural computations

As somatosensory information ascends from the periphery to cortex, spatiotemporal patterns of peripheral population activity are subject to spatial and temporal differentiation (variation) computations (Connor and Johnson 1992; DiCarlo and Johnson 2000; Saal et al. 2015). We built a simple model of these variation filters by combining the responses of peripheral afferents to textures presented at multiple speeds. Specifically, we first temporally smoothed afferent spiking responses using a filter designed to mimic an excitatory post-synaptic potential (EPSP) (Bengtsson et al. 2013):

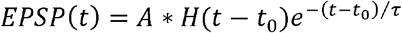

where *t*_0_ is the time of each spike, *τ* is a decay time set to 5 ms, *H*(*t*) is the Heaviside step function, and *A* is a normalization constant such that ∫ *dt EPSP*(*t*) = 1. The integration time of temporally lagged inhibition is generally longer than that of excitation (DiCarlo and Johnson 2000; Sripati et al. 2006), so for the temporal variation inhibitory field (see below), we computed the post-synaptic potentials with *τ* set to 9 ms.

This way, we generated a large population of temporally aligned post-synaptic potential traces, to which we applied spatial and temporal variation computations as described below. For each texture and speed, we first aligned all the trial-averaged traces in the population (between 21 and 39 neurons, depending on the speed). To this end, we first found the strongest cross-correlation between any two pairs of neurons (within a maximum shift of 300 ms), averaged them at their optimal lag, and then found the strongest cross-correlation between this new trace and any of the remaining traces. This procedure was repeated until the full population was aligned. As we only had one 2-second trace for any given texture at 40 mm/s, these traces were split in half and then maximally aligned to create two 1-second traces.

Next, we computed pseudo-populations of “downstream” neurons that implement both spatial and temporal variation filters. We implemented temporal variation by subtracting out a delayed version of the same spike train (this time smoothed over 9 ms), at random delays (sampled evenly between 15 and 50 ms) and weights (0 to 50% of the excitatory weight), the ranges of which were selected based on previously documented spatial-temporal receptive fields of cortical neurons (DiCarlo and Johnson 1999, 2000; Sripati et al. 2006; Saal et al. 2015). We implemented spatial variation by subtracting from one response trace the trace derived from a different, randomly selected afferent of the same type (SA1, RA, or PC). We simulated the relative spatial location of the inhibitory subfield as trailing (along the axis defined by the scanning direction) 2 to 4 mm (randomly selected) behind the excitatory subfield at a random weight (0 to 50% of the excitatory weight), again inspired by previous findings (DiCarlo and Johnson 1999, 2000; Sripati et al. 2006). In practice, this meant subtracting out the spatial inhibition at a temporal delay that shifted for different speeds. After subtracting out the temporal and spatial inhibition, we half-wave rectified each trace and summed it to obtain the “output” of that simulated neuron. This procedure was repeated for two trials of every texture response at every speed, and then repeated 100 times for each afferent, each time with a different set of randomized parameters. From the responses of each afferent, then, we simulated the responses of a set of neurons that each reflected an idiosyncratic variation computation.

We then computed the correlation between each simulated cortical response and the simulated response averaged across the population (24 shared textures, 80 mm/s, all 141 neurons). We also computed the speed/texture ratio from the simulated responses of each neuron (as defined above). The median correlation reported in the text and the cumulative distribution in *Figure 4A* were computed across all permutations and afferents. To compute the alignment index for the simulated population, we first randomly selected a simulated neuron derived from each of the 21 afferents. We then computed the alignment index on this pseudo-population as described above. We repeated this procedure over 200 random selections of simulated neurons.

## RESULTS

We have previously reported the texture-evoked responses of 39 tactile afferents from six anesthetized macaque monkeys (Weber et al. 2013) and 141 neurons from the somatosensory cortices of three awake macaque monkeys (Lieber and Bensmaia 2019), as many different textures were scanned over the skin. For a subset of these neurons, we were able to maintain isolation quality long enough to run a second protocol of textures presented at multiple speeds. This protocol was run on 21 tactile fibers – 9 slowly adapting type 1 (SA1), 9 rapidly adapting (RA), and 3 Pacinian (PC) fibers — and 49 neurons in somatosensory cortex – 14 from Brodmann’s area 3b, 26 from area 1, and 9 from area 2 – with receptive fields on the distal fingertip. For the peripheral nerve experiments, each of 55 different textures was scanned over the skin at 3 different speeds (40, 80, and 120 mm/s) using a rotating drum stimulator. For the cortical experiments, each of 10 different textures was scanned over the skin at 4 different speeds (60, 80, 100, and 120 mm/s), which spans the range of speeds spontaneously used to explore tactile textures (Morley et al. 1983; Gamzu and Ahissar 2001; Libouton et al. 2010; Tanaka et al. 2014; Callier et al. 2015). We sought to determine the effect of scanning speed on the neural representation of texture, and how these representations change between periphery and cortex.

### Texture responses are modulated by scanning speed

We found that, for both tactile nerve fibers and cortical neurons, increasing the speed at which a texture is scanned across the skin drives an increase in the firing rate response (Figure *1A-C*). This effect was significant for nearly every tactile nerve fiber (20/21, *p* <0.05, permutation test) and for a majority of neurons in cortex (37/49). For consistency, we only included in the analysis the responses to textures that were either shared across the peripheral and cortical experiments or were similar (Supplementary Figure *1A*, see Methods). Using this matched set of textures, we compared the speed-sensitivity of peripheral afferents to that of cortical neurons by expressing sensitivity as a percentage increase in firing rate per doubling in speed (Supplementary Figure 1*B-D*). Using this metric, we found that peripheral afferents exhibited more speed-sensitivity than did cortical neurons (Figure *1D*)(median across cells ± median absolute deviation: periphery: 28.7%±5.1%, cortex: 20.0%±11.5%, *p*<0.05 Wilcoxon rank-sum test; also see Supplementary Figure *1E* and Supplementary Table *1*). This difference was almost entirely driven by a subpopulation of cortical cells with highly speed-invariant responses (13/49 with speed-sensitivity < 10% increase/doubling, 0/21 peripheral afferents < 10%). While we did observe significant differences in speed-sensitivity across different afferent classes (Delhaye et al. 2019), every submodality trended towards more speed-sensitivity than that was observed in cortex (RA: 29.3%±3.3% and PC: 35.0%±5.4%, p < 0.05, SA1: 23.8%±6.2%, p=0.18, see *Supplementary Figure 2A-B*). The speed-sensitivity of responses in area 3b trended towards smaller values than responses from areas 1 and 2 (area 3b: 12.9%±9.8% < area 1: 23.1%±7.5% at *p*<0.05 and area 2: 24.4%±24.9%, *p*=0.33, see Supplementary Figure 2A-B), consistent with previous reports (Dépeault et al. 2013; Bourgeon et al. 2016).

**Figure 1.**
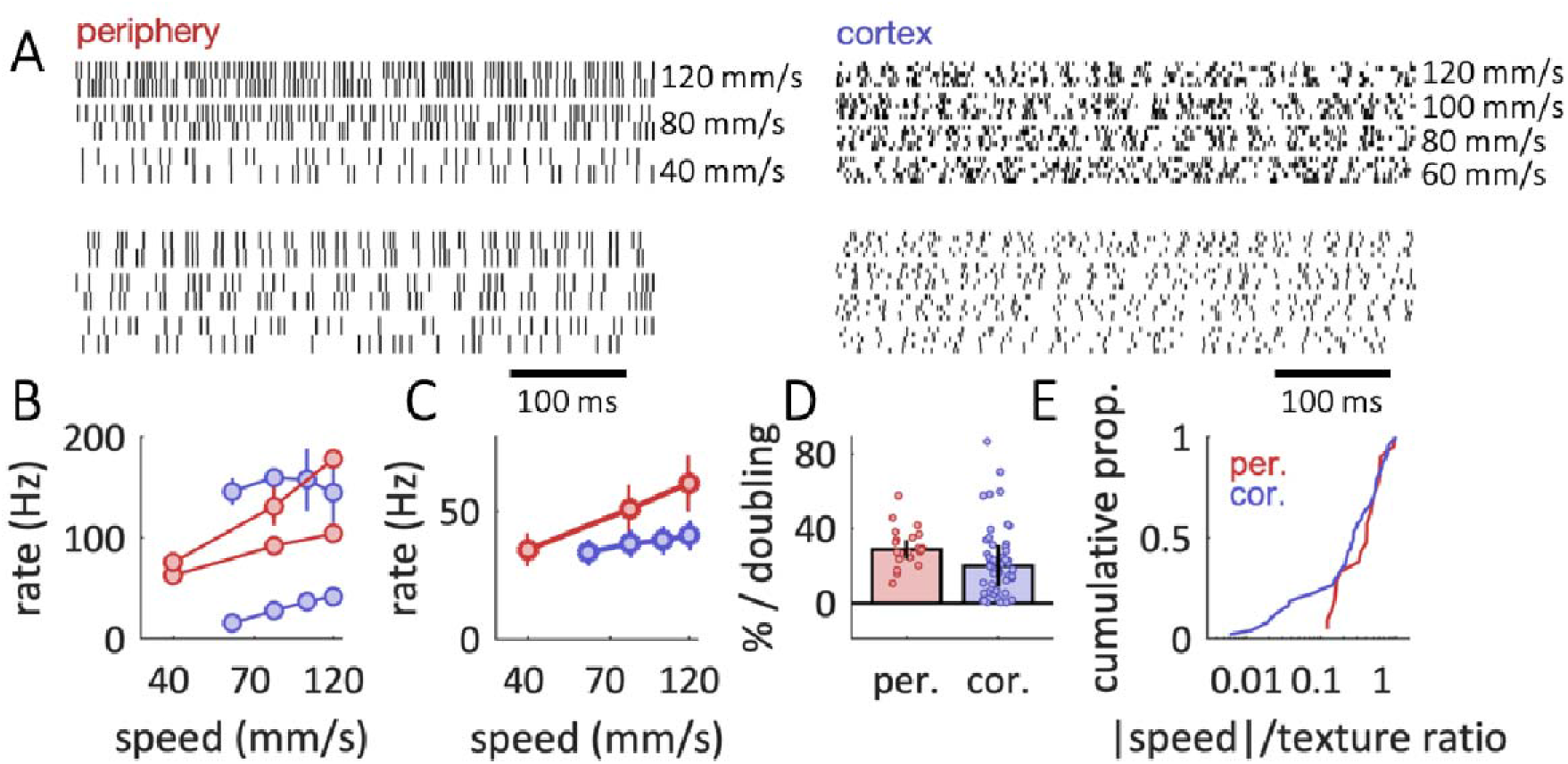
Texture responses are modulated by scanning speed. A| Spiking responses from two example tactile nerve fibers (top: RA, bottom: PC) and two cortical neurons (top: area 3b, bottom: area 1) to a texture (hucktowel) presented at three/four different speeds, respectively. B| Average firing rate of the spiking response shown in A| for the two afferents (red) and two cortical neurons (blue). Speeds on the abscissa are plotted on a log scale. Error bars are standard deviations across trials. C| Average firing rate across all textures and cells at the periphery (red) and in cortex (blue) vs. scanning speed. Speeds on the abscissa are plotted on a log scale. Error bars denote standard errors across cells, textures, and trials. D Median speed effect, reported as the percentage increase in mean population firing rate per doubling in speed (normalized to the firing rate at 80 mm/s). Different points denote different neurons. Error bars are median absolute deviations across cells. E| Cumulative distribution of the ratio between the speed effect and texture effect for peripheral afferents (red) and cortical neurons (blue). While the medians of the distributions are similar, the distribution derived from cortex contains a large proportion of neurons whose responses to texture are strongly speed-independent.

Next, we assessed whether the decreased speed-sensitivity in cortex resulted in a more speed-invariant representation of texture. To this end, we computed the ratio between the texture-dependence of firing rates and their speed dependence (Figure *1E*). The resulting speed/texture ratio, lower for more speed invariant texture coding – was only marginally lower in cortex than in the periphery (median ratio of speed to texture-sensitivity, periphery: 0.59, cortex: 0.47, *p*=0.287 Wilcoxon rank-sum test), suggesting that cortical texture representations are only slightly more speed invariant in cortex than at the periphery. Indeed, while speed-sensitivity tends to decrease in cortex, so does texture-sensitivity. To directly test the ability of individual neurons to discriminate between textures, we computed a signal-to-noise ratio (SNR) as a metric of discriminability for texture pairs (see Methods). We found that, for any given speed difference, SNR values were largely similar for individual peripheral and cortical neurons (*Supplementary Figure 3A*). Thus, at the single-cell level, speed had a largely similar effect on peripheral and cortical texture responses.

Given that median speed-sensitivity was similar between the peripheral and cortical populations, we next considered whether speed-invariance might be achieved by a specialized subpopulation of cortical neurons. As signals ascend the somatosensory hierarchy, the tuning of individual neurons becomes increasingly heterogeneous (Lieber and Bensmaia 2019). We might thus expect subpopulations of cortical neurons to show specialization for speed or texture coding (Dépeault et al. 2013; Bourgeon et al. 2016). Indeed, we found a significant proportion of cortical cells exhibited speed/texture ratios weaker than any observed in peripheral afferents (Figure *1E*) (12/49 with ratio < 0.12, p<0.05, permutation test, see Methods), an effect that was present in all three cortical fields (*Supplementary Figure 2C*). Therefore, somatosensory processing does not simply extinguish speed-sensitivity as signals ascend from the periphery to somatosensory cortex but rather creates a wide range of response properties in cortex that could potentially represent information about both texture and scanning speed.

### Texture and speed signals are more independent in cortex than at the periphery

The diverse tuning of individual cortical neurons suggests that the population representations of texture and speed may be more independent in cortex than at the periphery. To this end, we first identified linear combinations of neurons within each population that best captured either texture-driven or speed-driven modulations of firing rate.

To identify the most texture-sensitive dimensions in the peripheral and cortical populations, we applied principal components analysis (PCA) to each population’s response to 24 textures presented at a single speed (80 mm/s). The PCA yielded principal components [PCs], ordered by their ability to account for variance in the texture response. To identify the speed-sensitive dimension in each neural representation, we applied demixed principal components analysis (dPCA) (Kobak et al. 2016) to each population’s response to 10 textures presented at multiple speeds. Although this analysis identified multiple speed-sensitive dimensions in each population response, a single dimension captured most of the speed-related response variance (peripheral: 87.6%, cortical: 63.9%, see Methods) and significantly tracked speed magnitude across different textures (*R*^2^ to log_2_ speed, periphery: 0.64, cortex: 0.24, *p* < 0.01, F-test).

To examine the relationship between the texture and speed representations, we assessed the extent to which the texture- and speed subspaces were orthogonal. That is, to what degree do changes in speed affect the speed representation but not the texture representation, and vice versa? To this end, we computed the proportion of speed-related variance that was captured by each dimension of the texture representation (*Figure 2C*), a quantity we refer to as the alignment index (Elsayed et al. 2016; Gallego et al. 2018)(see Methods). We found that speed-driven changes were primarily captured by the first principal component of the texture representation and this relationship was far stronger in the periphery than in cortex (average alignment index, peripheral: 0.80, cortical: 0.46). This effect was robust across the full texture space (*Figure 2D*), where the peripheral representations of texture and speed were still more closely aligned than were their cortical counterparts. Surprisingly, this effect was not simply a consequence of a subpopulation of particularly speed-invariant neurons. Rather, we found that the increased separation of speed and texture was robustly present even when the most speed-invariant cortical neurons were removed (*Supplementary Figure 4*). We surmised that the increased independence of the cortical texture response could endow it with an increased ability to support speed-independent texture discrimination. We extended our SNR analysis to populations of neurons (see Methods) and indeed found that cortical subpopulations showed stronger discriminability of texture pairs than comparably sized populations of tactile fibers (*Supplementary Figure 3B-C*), a strong contrast to the largely similar performance seen for individual cortical neurons and afferents. We conclude that, as texture-driven responses ascend the somatosensory hierarchy, populations of neurons encode speed and texture information in increasingly independent representations that support speed-invariant texture discrimination.

**Figure 2.**
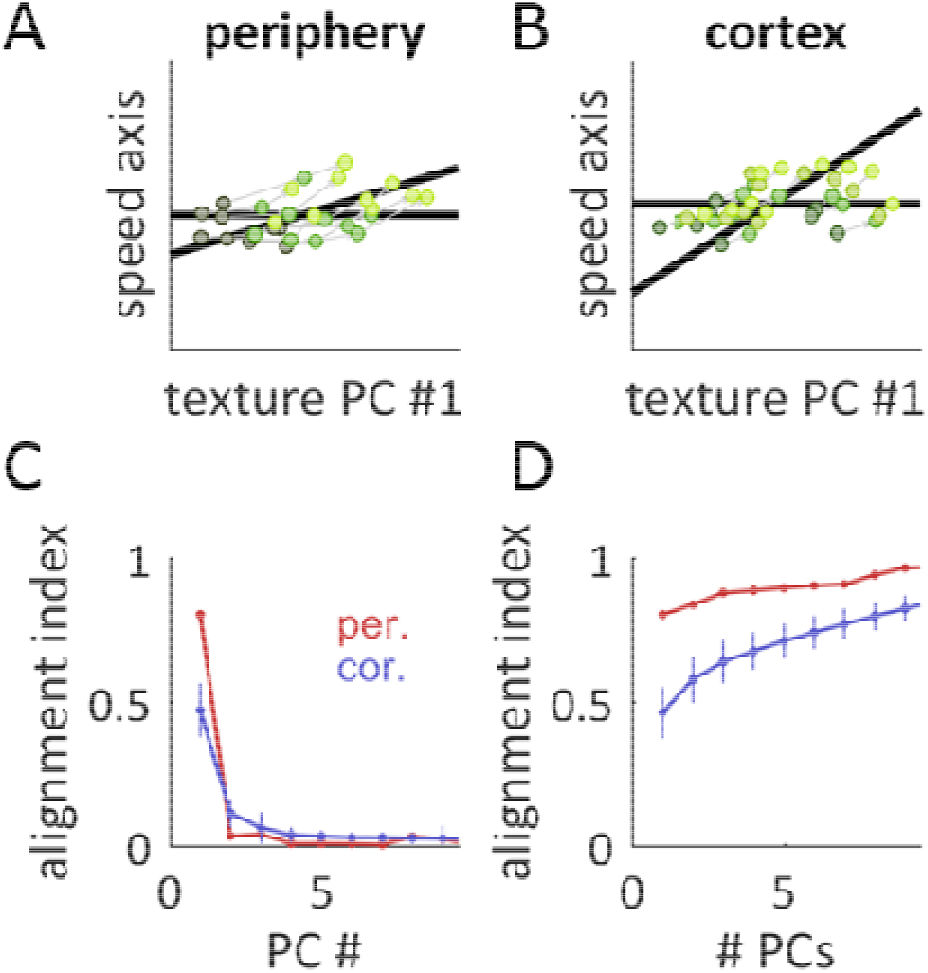
Texture and speed signals are more independent in cortex than at the periphery. A-B| Texture population responses are projected to a two-dimensional plane. One dimension (plotted on the abscissa, and also as a horizontal black line) corresponds to the first principal component of the texture space (found by applying PCA to texture responses at 80 mm/s). A second dimension (corresponding to the oblique black line) corresponds to the primary speed axis (found by applying dPCA to texture responses at multiple speeds). The primary texture and speed axes are not orthogonal to each other and a more acute angle between the two black lines indicates greater overlap in the representations of texture and speed. Accordingly, the ordinates of the plots correspond to the component of the speed axis orthogonal to the primary texture axis. Texture responses are plotted at three speeds (principal components for 10 different textures plotted at different speeds, low speeds: dark green, high speeds: light green). Speed-driven changes in firing rate are less aligned to the primary texture axis in corte than at the periphery. C| Alignment index between the primary speed axis and individual texture axes for the peripheral (red) and cortical (blue) populations. The alignment index measures the proportion of speed-related variance captured by each texture dimension (see Methods). Error bars for the cortical population (blue) are the standard deviation across randomly sampled subpopulations of 21 neurons. D| Alignment index between the primary speed axis and the multidimensional texture space for the peripheral (red) and cortical (blue) populations, shown for the first 9 principal components of the texture space.

### Cortical responses account for speed-invariant texture perception

Next, we examined the extent to which neuronal responses could account for the well-documented speed-invariance of texture perception (Lederman 1974; Meftah et al. 2000; Boundy-Singer et al. 2017). To this end, we tested the hypothesis that perceived roughness is determined by the population firing rate in somatosensory cortex (Burton and Sinclair 1994; Lieber and Bensmaia 2019) using a previously published set of roughness ratings obtained from human subjects. First, we regressed roughness ratings onto the population firing rate evoked when textures are scanned across the skin at 80 mm/s (*Figure 3A*)(cross validated *R*^2^, peripheral: 0.81, cortical: 0.77). Next, we assessed how well this linear model could account for the neuronal responses at other speeds (*Figure 3B*). We found that roughness estimated from both peripheral and cortical responses were strongly modulated by changes in scannin speed (% increase in roughness per doubling of speed, periphery: 29.9%, cortex: 19.4%) in contrast to the roughness ratings, which were largely speed-independent. Given the observed heterogeneity of cortical tuning, we next considered that the neural code for roughness might rely more on some neurons than others (Chapman et al. 2002; Bourgeon et al. 2016). To test this hypothesis, we regressed the first five texture-related principal components of each population response on perceived roughness (*Figure 3A-B*, middle bars). This led to a more accurate prediction of perceived roughness (cross-validated *R*^2^, peripheral: 0.87, cortical: 0.81), but only marginally reduced the speed dependence of the roughness predictions (% increase in roughness per doubling of speed, periphery: 25.8%, cortex: 15.3%).

**Figure 3.**
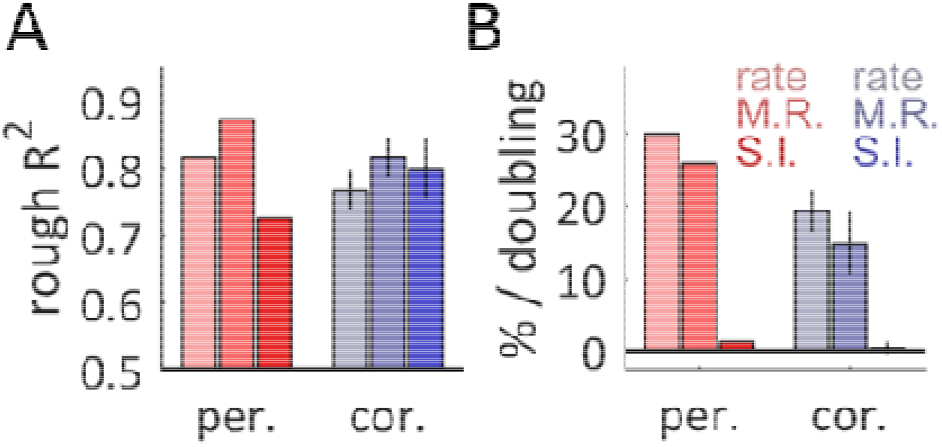
Cortical responses account for speed-invariant texture perception. A| Cross-validated *R*^2^ between the predicted and true roughness, for predictions based on peripheral (red) and cortical (blue) population responses (populations of 21 cells). From left to right, bars represent predictions based on regressions of the population averaged firing rate (light, rate), the best-fit regression of 5 principal components (medium, M.R.), and a best-fit regression constrained to minimize speed dependence (dark, S.I.). Error bars denote standard deviations across different samples of 21 cells. B| Speed dependence of the roughness prediction reported as a percentage increase in firing rate per doubling of speed, with color conventions as in A| The cortical population can support a speed-independent prediction of roughness, while the peripheral population cannot simultaneously support speed-independence and an accurate roughness prediction.

To create fully speed-independent roughness predictions, we next constrained our regression weights to be orthogonal to the primary speed-related dimension (found using dPCA) in each population (*Figure 3A-B*, right bars). This approach successfully eliminated the speed-dependence for both sets of roughness predictions (% increase in roughness per doubling of speed, periphery: 1.2%, cortex: 0.2%). However, for the peripheral population, enforcing speed-independence strongly reduced the predictive power of the peripheral model (peripheral cross-validated *R*^2^: 0.73, from 0.87). In contrast, enforcing speed independence had essentially no effect on the predictive power of the cortical model (cortical cross-validated *R*^2^: 0.79 from 0.81). This effect was robust across a wide range of subpopulation sizes and regression parameters (*Supplementary Figure 5A-B*) but was highly variable for different individual subpopulations of peripheral afferents (*Supplementary Figure 5C*). That is, while some subpopulations of afferents reached levels of roughness prediction that nearly matched their cortical counterparts, others failed catastrophically. Thus, the cortical population response contains a robust, speed-independent readout of perceived roughness that is not present in the peripheral firing rate response.

### Known cortical computations account for the untangling of speed and texture information

As information ascends any sensory neuraxis, neural representations are repeatedly transformed by a set of canonical computations that shape the feature selectivity of downstream neurons. One well-established transformation in the somatosensory system is the computation of spatial (Connor and Johnson 1992; DiCarlo and Johnson 2000; Lieber and Bensmaia 2019) and temporal (Saal et al. 2015) variation: the extent to which the peripheral neural representation exhibits inhomogeneous (“edge-like”) structure in space or time. We hypothesized that these differentiation computations could also give rise to an increasingly heterogeneous population response to texture and speed as signals ascend from periphery through cortex. Indeed, increased response heterogeneity has been proposed as an organizing principle for the structure of receptive fields in the visual and auditory systems (Olshausen and Field 1996, 2004; Van Hateren and Ruderman 1998; Lewicki 2002). To test this hypothesis, we built a neurally plausible model of spatial and temporal variation using the responses of peripheral afferents. Specifically, we added two subtractive influences to each modeled cell: spatially offset inhibition originating from a separate afferent, and temporally offset inhibition tracking the cell’s response but with the sign inverted (Figure *4A*). Using a range of biologically plausible model parameters (see Methods), we investigated whether pseudo-populations of such simulated neurons exhibited speed-invariant texture coding.

**Figure 4.**
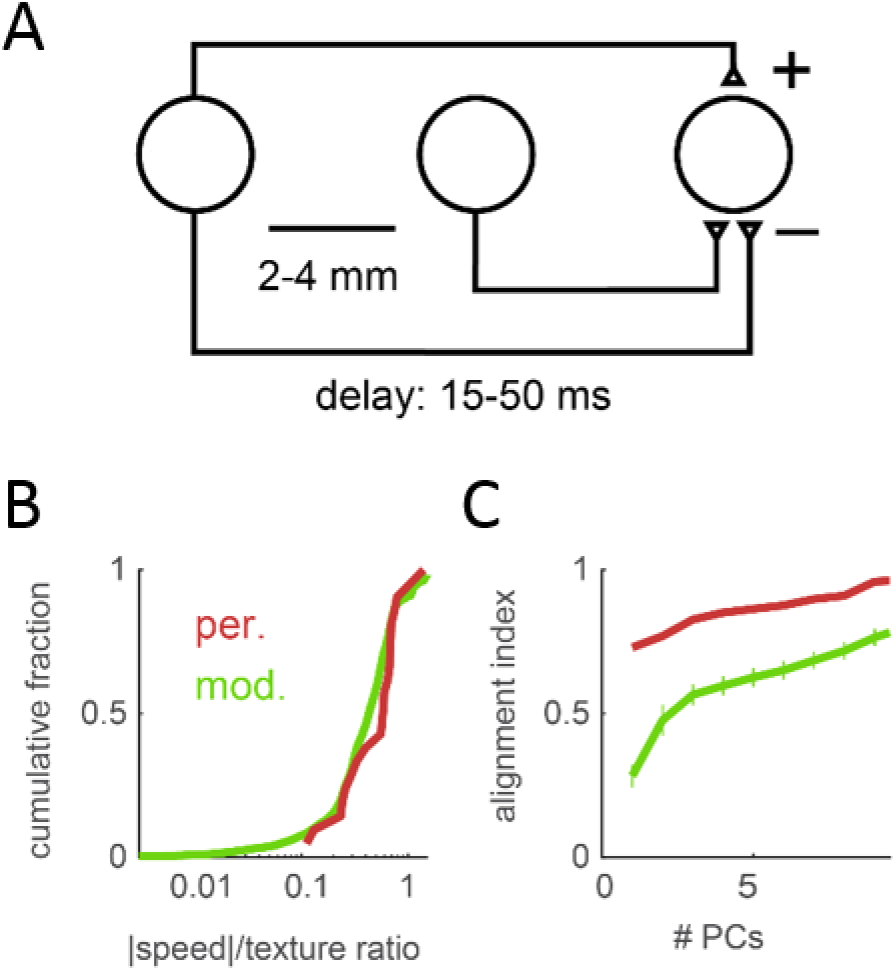
Known cortical computations account for the untangling of speed and texture information. A| Cartoon of the variation model. The texture response of a single afferent is combined with a spatial variation signal, modeled as an inhibitory response stemming from an afferent whose receptive field is located 2 to 4 mm away along the scanning direction, and a temporal variation signal, modeled as an inhibitory copy of the original afferent’s response delayed by 15-50 ms. B| Cumulative distribution of the ratio between the speed effect and texture effect for tactile fibers (red) and simulated cortical neurons (green). C| Alignment index (as in Figure *2*) between the speed and texture spaces for small populations (*N*=21 units) of tactile fibers (red) and simulated neurons (green), shown for the first 9 principal components of the texture space. Error bars denote the standard deviation across subsamples of simulated responses. The variation model exhibits a greater separation of speed and texture information than does the afferent population.

We first verified that the outputs of the variation model resembled actual cortical responses to texture (median correlation with population-averaged cortical response to 24 textures scanned at 80 mm/s: variation model outputs *r*=0.71, individual cortical neurons *r*=0.75). Next, we evaluated the speed-sensitivity of individual “downstream” model outputs by calculating their speed/texture sensitivity ratio (as above). Outputs of the variation model showed comparable levels of speed-sensitivity to peripheral afferents (*Figure 4B*)(speed/texture ratio, median ± median absolute deviation, model: 0.43±0.18 vs. afferents: 0.59±0.18). However, when we examined population-level representations in model outputs using the texture/speed alignment index (as above), we found that small populations (N=21 units) separated texture and speed more robustly than did peripheral afferents (*Figure 4C*). Comparing these results to models built with only spatially or temporally offset inhibition, we found that this separation was most strongly driven by spatial variation mechanisms, although temporal variation mechanisms contributed significantly to the separation as well (Supplementary Figure *6*). Thus, our simulation suggests that the well-established cortical computations of spatial and temporal differentiation contribute to the heterogeneous speed- and texture-sensitivity observed in somatosensory cortex, which in turn underlies the untangling of speed and texture representations.

## DISCUSSION

The perception of texture is remarkably tolerant to changes in scanning speed. Indeed, psychophysical ratings along the three principal perceptual axes of textures – roughness, hardness, and stickiness – are identical across speeds (Lederman 1974; Meftah et al. 2000; Boundy-Singer et al. 2017). Furthermore, the perceived dissimilarity of a pair of textures – which probes texture perception across all of its dimensions and attributes – is very similar whether the two textures are scanned at the same speed or at different speeds (Boundy-Singer et al. 2017). What makes this perceptual invariance so remarkable is that the response of tactile nerve fibers are highly speed dependent. Indeed, in the nerve, texture-elicited firing rates increase with scanning speed, and temporal spiking sequences, which carry critical texture information, also change with speed. The central result of the present study is that the cortical population response exhibits a capacity for speed-invariant coding of texture that exceeds the capabilities of the peripheral population response. Therefore, somatosensory processing between the periphery and cortex yields overlaid representations of texture and speed that are relatively independent, and thus much easier to decode independently.

### Experimental caveats

The textures used in the peripheral and cortical experiments were designed to address different scientific questions and only overlap partially. To overcome differences in texture set, we analyzed the neuronal responses to a subset of textures that drives similar patterns of responses in somatosensory cortex (Supplementary Figure *1*A). The speeds used in the cortical experiments spanned a different range than did those in the peripheral experiments to better span the behaviorally relevant range (Callier et al. 2015). To overcome differences in speed range, we selected a metric of speed-sensitivity (change in firing rate per doubling in speed) that is consistent at low and high ranges of speed (see Methods). These features of the analysis address biases based on stimulation paradigm (textures, speeds) that might skew our results. Note that we reach the same conclusions when using the full set of textures and speeds (Supplementary Figure *1*E) and that our measurements of peripheral and cortical speed-sensitivity are consistent with those observed by other groups (Supplementary Table *1*).

In the present study, we use neural responses measured in macaques to predict texture perception in a different species: humans. Human and macaque hands show very similar patterns of cutaneous innervation (Johansson and Vallbo 1979; Darian-Smith and Kenins 1980; Paré et al. 2003) and the tactile nerve fibers in the two species exhibit nearly identical response properties (Johansson et al. 1982; Phillips et al. 1992). Macaques can successfully perform texture discrimination tasks (Chapman and Ageranioti-Bélanger 1991; Tremblay et al. 1996), and human texture perception can be successfully predicted by the responses of macaque nerve fibers (Connor et al. 1990; Connor and Johnson 1992; Blake et al. 1997; Weber et al. 2013; Lieber et al. 2017) and cortical neurons (Bourgeon et al. 2016; Lieber and Bensmaia 2019). We therefore believe that the macaque model of texture and speed coding is well suited to account for human texture perception.

### Previous work on the speed invariance of texture representations

We find a continuum of response properties in somatosensory cortex, from neurons that are as sensitive to speed as are peripheral afferents to neurons that are nearly speed-independent. These data are broadly consistent with previous studies that have emphasized that different cortical neurons show responses that are purely texture-selective, purely speed-selective, or responsive to both texture and speed (Tremblay et al. 1996; Dépeault et al. 2013; Bourgeon et al. 2016). However, we emphasize that there exists significant response heterogeneity within these subpopulations as well, and that the distribution of speed-sensitivity across cortex is likely better described as a continuum than as a bimodal distribution of texture and speed specialists.

### Variation computations give rise to speed-invariant representations of texture

In the above analyses, we compare texture representations in afferent firing rates and cortical firing rates. However, a large body of evidence suggests that stimulus information is not encoded simply in the firing rates of tactile nerve fibers, but rather in spatio-temporal patterns of activation (Talbot et al. 1968; LaMotte and Mountcastle 1975; Connor and Johnson 1992; DiCarlo and Johnson 2000; Mackevicius et al. 2012; Weber et al. 2013; Birznieks and Vickery 2017). These putative peripheral neural codes imply computations along the neuraxis where spatio-temporal motifs in the afferent input are converted to firing rate codes downstream. One of the canonical computations is that of differentiation, both spatial and temporal. Indeed, neurons in somatosensory cortex have been shown to exhibit Gabor-like spatial receptive fields, reflecting a spatial differentiation (DiCarlo and Johnson 2000; Sripati et al. 2006; Bensmaia et al. 2008), and bi-lobed temporal receptive fields, reflecting temporal differentiation (DiCarlo and Johnson 2000; Sripati et al. 2006; Saal et al. 2015). We demonstrate that these computations confer an additional benefit to the cortical code for texture: increased separability of information about texture and speed. This separability likely reflects a broader sensory function, namely to create a basis set that efficiently and sparsely encodes behaviorally relevant features across the breadth of naturally occurring stimuli (Olshausen and Field 2004). In the somatosensory system, Gabor-like spatial and temporal filters transform a largely homogeneous set of peripheral responses into a widely divergent set of cortical responses, as has also been shown in the visual system (Olshausen and Field 1996; Van Hateren and Ruderman 1998).

We emphasize that a single variation filter, by itself, is not sufficient to create response heterogeneity in a neural representation. Consider the receptive field structure of neurons in the lateral geniculate nucleus (LGN). Although these neurons do compute spatial variation, they do so using a spatial receptive field structure (center-surround) that is relatively homogeneous within any local population (Derrington and Lennie 1984). One synapse later, local populations of neurons in primary visual cortex exhibit receptive field structures that vary widely in their spatial extent, spatial frequency, and orientation (De Valois et al. 1982). This expansion in response properties is likely due to the increase in neural representation size between LGN and cortex. A comparable expansion exists between the somatosensory periphery and cortex, an important prerequisite for the observed heterogeneity in cortical responses. As such, we would attribute the separation of speed and texture information not just to the presence of variation filters per se, but rather to the wide range of different variation filters implemented along the somatosensory neuraxis. As inhibitory “variation-like” computations have been observed in the cuneate nucleus and ventral posterior thalamus (Bystrzycka et al. 1977; Lee et al. 1994), this separation of speed and texture information may begin to arise at earlier stages of processing and continue to progress at later ones.

### Invariance as a canonical sensory computation

To produce a stable percept of object identity, the somatosensory system must correct for the influence of speed from the texture representation. However, information about tactile speed is behaviorally relevant, and thus ideally would be preserved rather than eliminated. We find that the somatosensory system does not discard speed information but rather partitions texture and speed representations into increasingly separated subspaces. This partitioning is not perfect: the representation of tactile speed in somatosensory cortex is highly contaminated by texture identity and this influence of texture leads to a predictably non-veridical perception of tactile speed (Delhaye et al. 2019). Nonetheless, the preservation of any speed information speaks to the capacity of the cortical population representation to simultaneously encode many relevant variables simultaneously.

We find that, at the somatosensory periphery, object information (texture) and information about exploratory parameters (speed) are inextricably tangled together. As this mixed signal ascends the somatosensory hierarchy, it is transformed into two independently readable representations of these two parameters. This mirrors results from vision and audition, where successive levels of processing lead to a higher fidelity readout of both object identity (Rust and DiCarlo 2010; Town et al. 2018) and exploratory parameters (Hong et al. 2016). These convergent results suggest that separating (or untangling) information about objects and exploratory parameters is a canonical sensory computation (DiCarlo and Cox 2007).

## SUPPLEMENTARY FIGURES

**Supplementary Figure 1.**
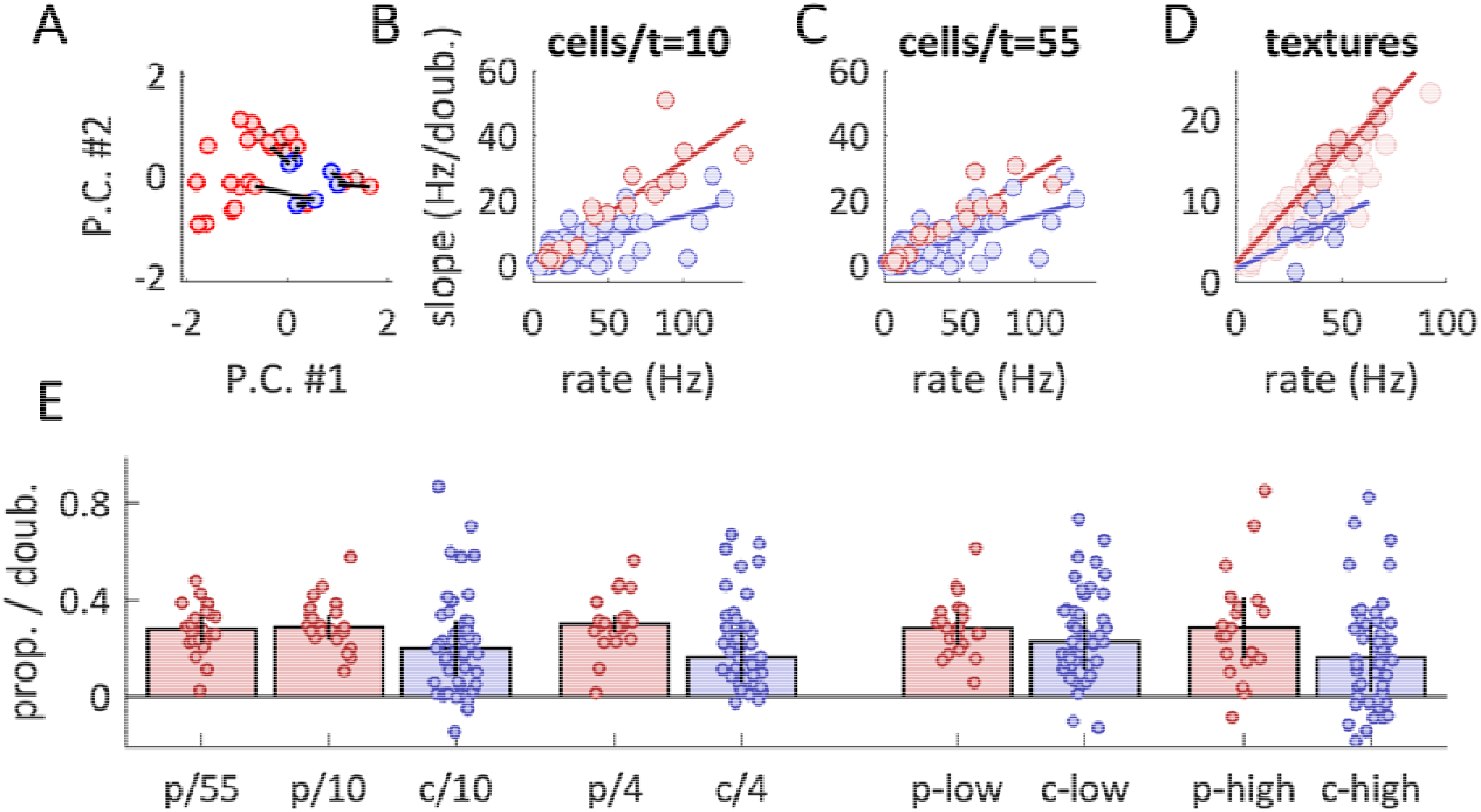
Different subsets of textures yield similar conclusions about speed-sensitivity. A| Cortical population responses to 30 different textures were projected onto the first two principal components of the texture representation (all presented at 80 mm/s). Four (4) of these textures were included in the speed set for both experiments (purple), twenty textures were included in the peripheral speed set but not the cortical one (red), and 6 textures (blue) were included in the cortical speed set but not the peripheral one. To fairly compare analyses periphery to cortex, we found the 6 textures from the peripheral speed set that evoked cortical responses most similar to the 6 non-overlapping textures from the cortical speed set (black lines). This set of 6 textures (combined with the set of 4 overlapping textures) was used for all analyses. B| Speed-sensitivity of individual neurons (measured in Hz / doubling of speed) vs. that neuron’s mean firing rate across textures, for peripheral afferents (red, using the set of 10 matched textures) and cortical neurons (blue). Each point denotes a cell. Speed slopes increase proportionally to each neuron’s average excitability (F-test for regression fit, both *p*<10-6). C| Speed-sensitivity vs. mean firing rate across textures as in B|, but afferent responses are averaged over the full set of 55 textures (F-test for regression fit, both *p*<10-6). Results are similar regardless of the texture set used. D| Population speed-sensitivity vs. mean firing rate across neurons, for individual textures. Dark red points indicate the matched peripheral set of 10 textures, light red indicate the other 45 textures. While peripheral slopes are closely related to the averaged population firing rates (F-test for regression fit, *p*<10-3), cortical slopes show a much noisier relationship (*p*=0.19). Different points denote different textures. E| Median normalized speed-sensitivity for the peripheral and cortical population for different sets of textures and speeds. For the five bars on the left: peripheral slopes were computed over either the full set of 55 textures, the matched set of 10 textures, or the exactly overlapping set of 4 textures. Cortical slopes were computed over the full set of 10 textures or the overlapping set of 4 textures. For the four bars on the right: peripheral and cortical slopes were computed using only “low” speeds (periphery: 40 and 80 mm/s, cortical: 60 and 80 mm/s) or “high” speeds (peripheral and cortical: 80 and 120 mm/s). All four sets of slopes were computed over the matched set of 10 textures. Cortical slopes are consistently lower than peripheral slopes, regardless of the texture set or speed range used to compute them.

**Supplementary Figure 2.**
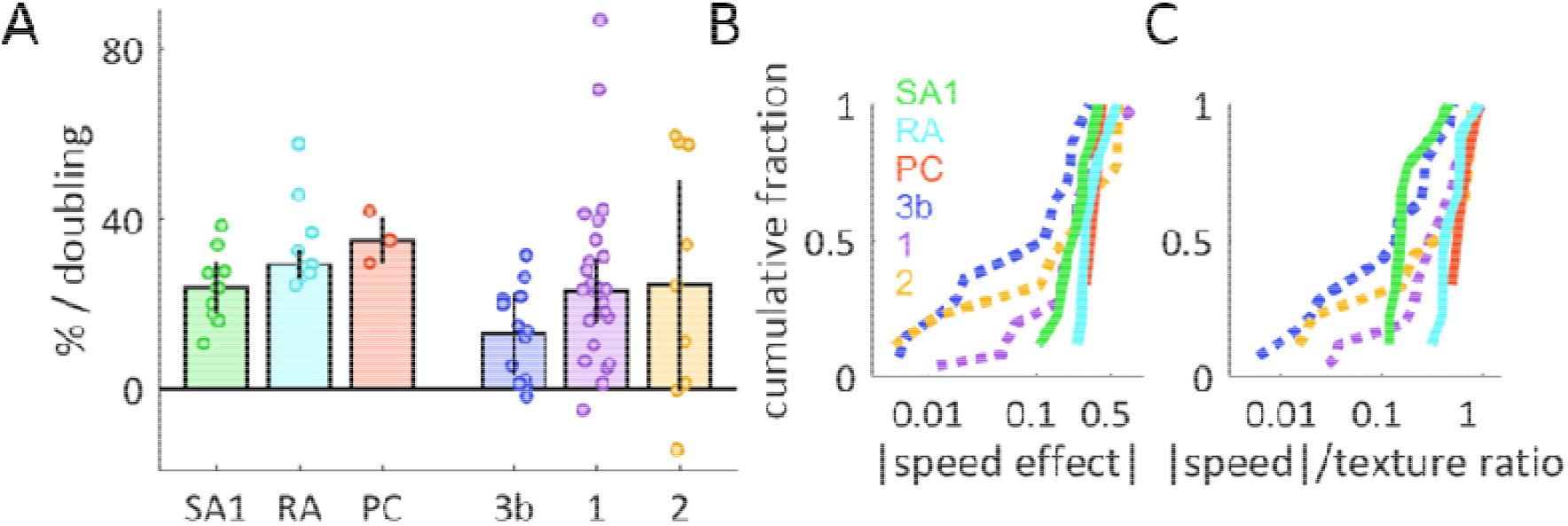
Speed-sensitivity differs across afferent classes and cortical fields. A| Speed-sensitivity for afferents and cortical neurons, broken out by afferent submodality and cortical field. Neurons in area 3b trend towards less speed-sensitivity than those in area 1 (*p*<0.05) or area 2 (*p*=0.33). B| Cumulative distribution of speed-sensitivity (absolute value) across afferents and neurons (same data as in A|, replotted). Colors represent afferent submodality and cortical area, as in A|. C| Cumulative distribution of the ratio between speed-sensitivity (as in B|) and texture coefficient of variation for each afferent and cortical neuron. All three cortical areas contain a significant subpopulation of neurons that exhibit stronger speed-invariance than the least speed-invariant SA1 afferent (fraction of neurons with a speed/texture ratio less than the lowest SA1 ratio: area 3b: 6/14, area 1: 4/26, area 2: 2/9, all *p* < 0.05).

**Supplementary Figure 3.**
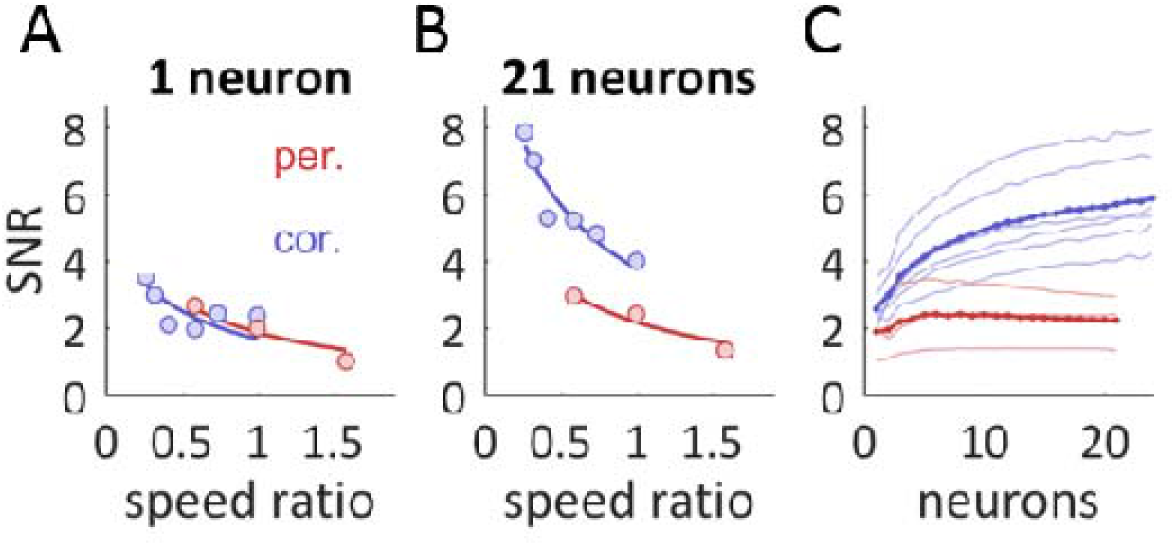
The cortical population representation of texture exhibits more invariance to changes in speed than does the peripheral representation. A| SNR vs. speed ratio, averaged across all texture pairs (10 textures) for peripheral afferents (red) or cortical neurons (blue). We defined SNR as the difference in response to each texture, averaged across a pair of speeds (the “signal”) divided by the spread of each texture’s firing rate across speeds (the “noise”). SNR was calculated over mean firing rates at two speeds which could be close together (left) or far apart (right). Best-fit curves of the form were fit to the data (see Methods). As expected, larger differences in speed drove lower discriminability for both tactile nerve fibers and cortical neurons, but discriminability is similar between the two populations at any given difference in speed. B| SNR vs. speed difference, now in multidimensional space for groups of 21 neurons. Discriminability was calculated along the neural dimension connecting the centers of the two textures. C| SNR vs. group size for individual conditions (light colors) and averaged across conditions (dark colors). Peripheral discriminability saturates quickly, while cortical discriminability grows rapidly for all conditions.

**Supplementary Figure 4.**
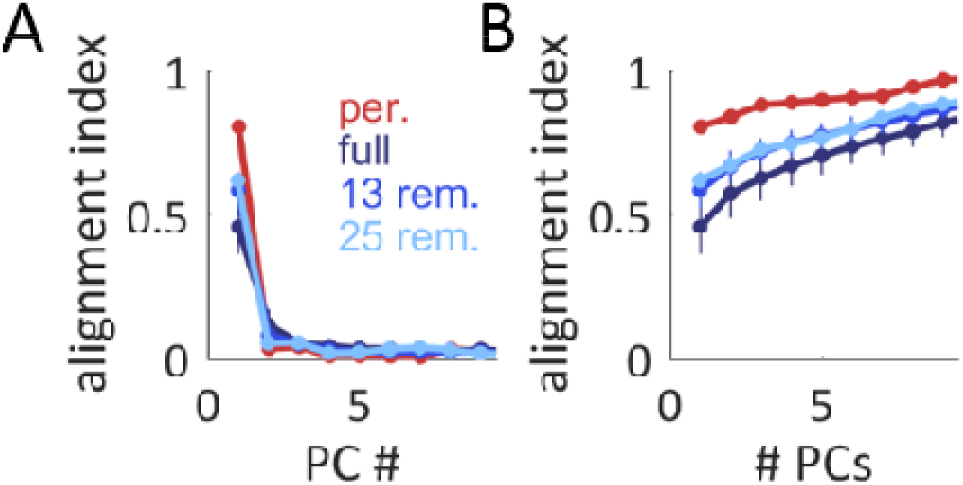
Cortical separation of texture and speed information is robust even among the most speed-sensitive subpopulations. A| Alignment index between the primary speed axis and individual texture axes, as in Figure 3C for three cortical subpopulations: the full cortical population (dark blue), a cortical population with the 13 most speed-invariant cells removed (medium blue, invariance measured using speed/texture ratio, as in Figure 2B), and a cortical population with the 25 most speed-invariant cells removed (light blue). B| Alignment index between the primary speed axis and the multidimensional texture space, as in Figure 3D. Colors as in A|. Even when the analysis is confined to the most speed-sensitive neurons, the cortical population response still shows more separation between speed and texture than does its peripheral counterpart.

**Supplementary Figure 5.**
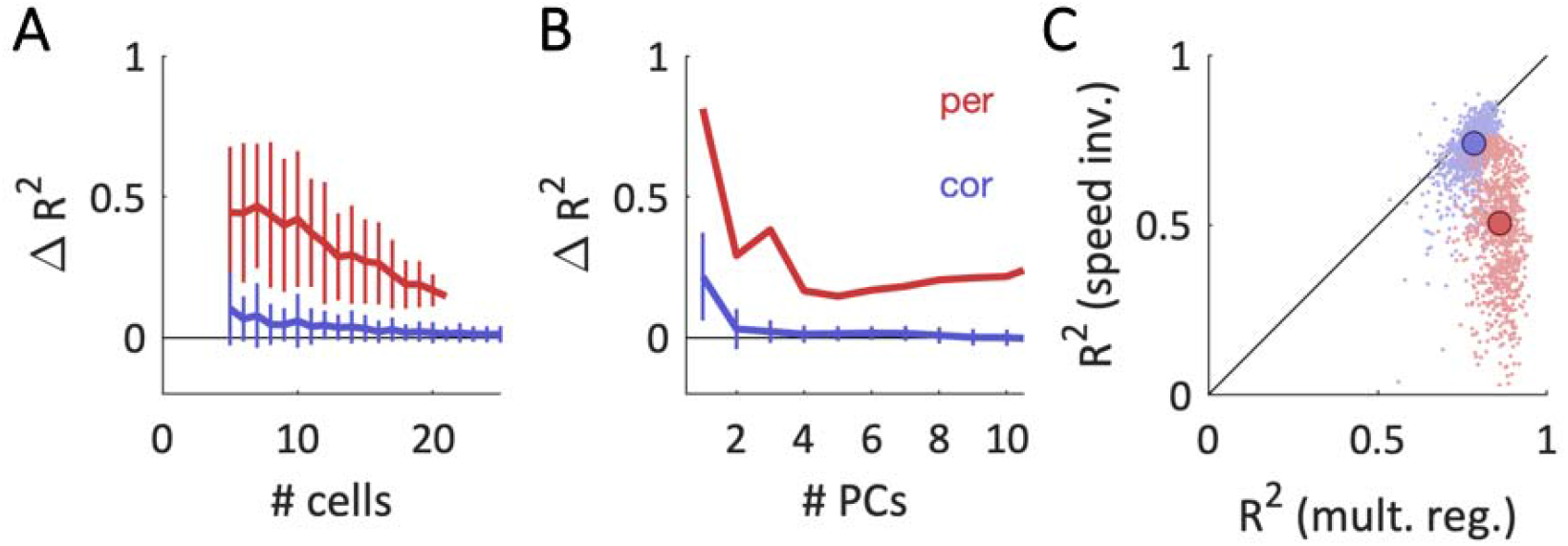
Predictions of perceived roughness are robust to changes in model parameters. A| Difference in cross-validated *R*^2^ between the full multiple regression model and the regression model constrained to be speed-independent, for neuronal subpopulations of different sizes (peripheral afferents: red, cortical neurons: blue). Each regression model was constrained to use the first 5 principal components of the population response – as such, we only plot results for subpopulations of size 5 and larger. B| Difference in cross-validated *R*^2^ between the two model classes, now for regression models trained using different numbers of principal components. All models were trained on subpopulations of 21 neurons. Cortical regression models were consistently more robust to the imposition of speed-independence. C| Performance of the speed-invariant model vs. performance of the full multiple regression model, for many different subpopulations of neurons (N = 10 cells), using 5 PCs. The speed-invariant performance of peripheral afferents was highly variable across subpopulations (median absolute deviation=0.12), compared to relatively stable performance of cortical subpopulations (median absolute deviation = 0.04). Despite this variability, peripheral subpopulations consistently exhibited larger drops in performance than cortical subpopulations (bigger peripheral drop for 96.8% of subpopulations). Thus, while some peripheral subpopulations showed cortical-level robustness, many catastrophically failed in their predictions of speed-invariant roughness.

**Supplementary Figure 6.**
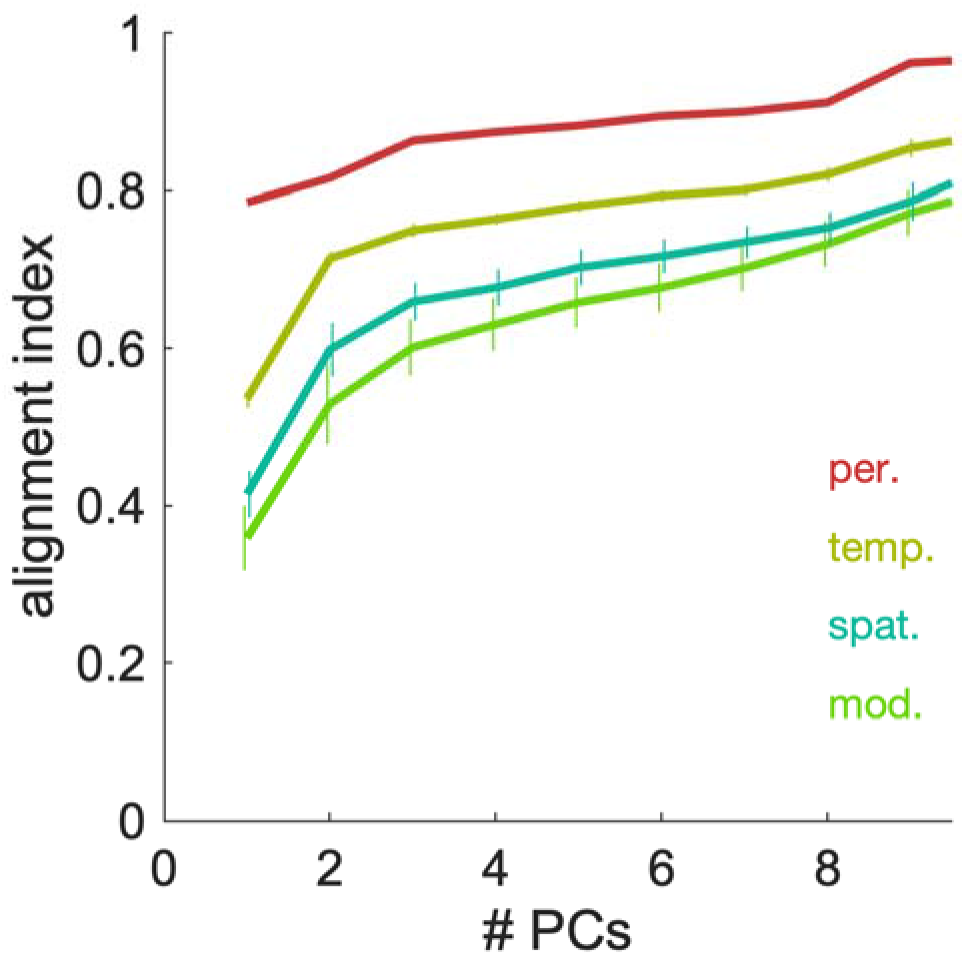
Both spatial and temporal mechanisms contribute to the separation of speed and texture information. Alignment index between speed and texture spaces for small groups of neurons (N=21) from four different populations. Indices for peripheral afferents (red) and full model outputs (green) are plotted as in Figure 4C. Also plotted are instantiations of the model that implement only feedforward inhibition (yellow, “temp.”) or surround inhibition (blue, “spat.”). Temporal or spatial variation, by themselves, can each increase the separability of speed and texture information. Models that implement both mechanisms exhibit less alignment than do those that only implement spatial variation (spatial and temporal < spatial alignment index for 95% of shuffled populations, for all # texture PCs conditions <= 6).

**Supplementary Table S1.** Previously published results on speed-sensitivity at the periphery and in cortex. Firing rate responses from previous studies were re-fit using the speed-sensitivity metric described in this paper (slope of the linear fit in log2(speed) coordinates). Results from (Goodwin and Morley 1987a) are based on the peak speeds of sinusoidal, back-and-forth sweeps, measured for a range of different periodic grating textures. Results from (Essick and Edin 1995) are in response to a brush sweeping over the skin, rather than a classically defined texture. Neural responses from (Sinclair and Burton 1991) were in response to macaque monkeys actively scanning textures with their finger (in contrast to the passive presentation of texture used in the other cited studies). As a result, the velocity and force used to scan the texture were under the volitional control of the animal, and thus 1) not consistent from trial to trial and 2) negatively correlated across trials. That is, unlike scans in the passive condition, scans at higher velocities tended to use less force. This may help explain the slightly lower levels of observed speed-sensitivity. Finally, the responses reported in (Dépeault et al. 2013) are not from the full population of somatosensory cortical neurons – rather, the authors only report the effect size of speed-modulation from the subset of neurons where that effect reaches statistical significance. This may help explain the slightly higher levels of observed speed-sensitivity. Overall, the levels of speed-sensitivity found in the present study broadly line up with those observed in previous work.

| Paper | Neural Subpopulation | Slope | Notes |
| --- | --- | --- | --- |
| Goodwin and Morley 1987a | SA1 | ~ 0% | Gratings, sinusoidal motion |
|  | RA | 23% | Gratings, sinusoidal motion |
|  | PC | 28% | Gratings, sinusoidal motion |
| Phillips and Johnson 1994 | Human SA1 | 24% | 2 mm spaced dot pattern |
|  | Human RA | 22% | 2 mm spaced dot pattern |
| Essick and Edin 1995 | Human, unspecified Aβ afferent | 14% | Brush stroked over RF |
| DiCarlo and Johnson 1999 | SA1 | 17% | Random dot pattern |
|  | RA | 30% | Random dot pattern |
|  | Cortical: 3b | 14% | Random dot pattern |
| Sinclair and Burton 1991 | Cortical: 3b & 1 | 14% | Actively scanned texture, scans at higher velocity were correlated with lower applied normal force |
| Dépeault et. al. 2013 | Cortical: 3b, 1, and 2 | 28% | Data from a subset of cortical neurons that exhibited statistically significant speed modulation |

## ACKNOWLEDGEMENTS

We would like to thank Alison Weber and Ju-Wen Cheng for collecting the peripheral nerve data, Zoe Boundy-Singer and Kristine Mclellan for collecting the psychophysical data, and Katie Long for helpful discussions. This work was supported by NINDS grant NS101325.

